# Th1 and Th2 cytokines profile of serum samples from dengue positive cases from Northeast India

**DOI:** 10.1101/2025.02.04.636426

**Authors:** Kiran Singh, Shinjini Bhattacharya, Ajanta Sharma, Sachin Kumar

**Author notes:** Corresponding authors: S. Kumar.

## Abstract

**Background:** Dengue fever is a fast-spreading acute arboviral infection transmitted by a human-Aedes mosquito cycle. Northeastern India is an endemic zone for dengue; regular outbreaks have been reported. However, molecular details of the virus infection in the host have not been established. Studies on dengue immunopathogenesis have primarily utilized data from American and Southeast Asian regions, with insufficient representation from India.

**Objective:** The present study aimed to characterize the Th1/Th2 cytokine expression profiles from dengue cases in different parts of northeastern India.

**Statistical analysis cum Results:** Among the panel of Th1/Th2 cytokines, IFN-γ, IL-2, IL-4, IL-5, IL-10, IL-13, and TNF-α cytokines are found to be higher in dengue-infected patients. Anti-inflammatory cytokines are somehow regulated differently in male-female patients.

**Conclusion:** This clinical study on dengue patient samples for cytokine profiling revealed distinct cytokine patterns and their regulation, but further investigation is needed to correlate with disease severity to identify potential biomarkers for disease progression.

## 1. Introduction

Mosquito-transmitted diseases like dengue considerably strain developing tropical countries, particularly those facing rapid urbanization, limited healthcare resources, and poor sanitation. [1, 2]. The causative agent dengue virus (DENV), is a member of the *Flavivirus* genus and family *Flaviviridae*. DENV is divided into four different genetically related serotypes but antigenically distinct [3].This disease is prevalent in tropical and subtropical regions and can pose life-threatening illness if left untreated. A complex interplay of environmental, social, and epidemiological factors shapes the geographical distribution of dengue. Globally, Latin America and the Caribbean, places like Brazil, Mexico, Colombia, and Cuba see high rates of dengue, primarily due to urban environments where mosquitoes can breed, especially in areas with poor water management [4, 5]. Moreover, continent-wise distribution of Dengue outbreaks has become a significant public health challenge. In Asia and the Pacific, countries like Thailand, Indonesia, the Philippines, and Vietnam often face frequent dengue outbreaks because of their warm, humid climates, perfect for *Aedes aegypti* mosquitoes to thrive. In India, the incidence of dengue has risen dramatically in recent decades, with major outbreaks reported in cities like Delhi, Kolkata, and Chennai. According to the reports of the National Centre for vector-borne diseases Control, there is a tremendous increase in the number of cases reported [6]. However, key factors such as rapid urbanization, inadequate sanitation, and climate change have contributed to the increased prevalence of the disease. [7-9]. Efforts to control mosquito vectors and improve surveillance remain critical in managing these outbreaks.

Northeast India’s geographical and climatic conditions play a significant role in the prevalence and transmission of dengue. The region has a tropical climate with high humidity and significant rainfall, particularly during the monsoon season, creating optimal breeding conditions for *Aedes* mosquitoes, the primary vectors of the dengue virus. The diversified landscape, which includes multiple water bodies and dense forests, contributes to mosquito multiplication. Furthermore, the region’s proximity to dengue-endemic nations like Bangladesh and Myanmar helps the virus spread across borders. Both urban and rural areas confront issues in mosquito control, with poor sanitation and infrastructure in some locations heightening the likelihood of outbreaks. Furthermore, population mobility between endemic areas raises the risk of dengue transmission. These variables highlight the importance of increased surveillance, vector management, and public health initiatives in reducing the effect of dengue in Northeast India.

The health concerns of different serotypes are vague and vary in severity, ranging from dengue fever (DF) to dengue hemorrhagic fever (DHF) to dengue shock syndrome (DSS) [10]. While primary infections are mild and primarily asymptomatic, secondary infections are more severe, including antibody-mediated enhancement (ADE) and cytokine storm. [11, 12]. Cytokines are signaling proteins released by immune cells in response to infections, and their levels can help differentiate between mild and severe forms of dengue. It has critical value in the immune response against flavivirus infections, such as Dengue, Zika, and Yellow Fever [13, 14]. Pro-inflammatory cytokines like IL-6, tumor necrosis factor (TNF-α), and IFN-γ are often elevated early, signaling viral infection. Dysregulated immune responses, including a cytokine storm, are linked to severe disease outcomes, such as hemorrhagic manifestations or shock. Cytokine profiles can also help distinguish between different flavivirus infections and predict the severity of the disease, assisting in timely diagnosis and treatment decisions. [15].

Additionally, specific cytokine patterns may offer insights into immune dysregulation and potential therapeutic targets [16]. A multitude of immunomodulatory molecules play a crucial role in developing dengue pathogenicity. Studying the function of cytokines in dengue pathogenesis is an intriguing topic. Dengue virus-infected cells produce several pro-inflammatory and anti-inflammatory cytokines, including IL-2, IL-4, IL-5, IL-6, IL-7, IL-8, IL-10, IL-12, IL-13, and TNF-α. [17-19]. Identifying pro-inflammatory and anti-inflammatory cytokines at the onset of symptoms may help predict the severity of dengue fever for treatment decisions.

This study evaluated Th1/Th2 specific cytokine profiles in dengue-positive patients’ serum samples. Studying differential cytokine expression in dengue patients will provide valuable information about immunity and disease mechanisms.

## 2. Study design

Blood samples were collected from 78 DENV-infected patients within 2-4 days of onset from different hospitals in Assam, India, 2024, with written informed consent (Table 1). All patients were further confirmed to have dengue by detecting the DENV antigen using the NS1 assay and DENV-specific antibodies through an in-house capture IgM Enzyme-Linked Immunosorbent Assay (ELISA) (NIV, ICMR, Pune, India). Blood serum samples were processed by spinning them at 1500 rpm for 10 minutes and then storing them at −80°C for future use. Levels of 9 different cytokines, including IL-2, IL-4, IL-5, IL-10, IL-12, IL-13, IFN-γ, granulocyte-macrophage colony-stimulating factor (GM-CSF), and TNF-α were assessed in the serum of patients and healthy subjects taken as control. These levels were analyzed using the Bio-Plex human cytokine Th1/Th2 9-plex panels (Bio-Rad Inc., Hercules, CA, USA). The process briefly involves mixing serum with beads coated with specific antibodies against target cytokines and then incubating with biotinylated anti-cytokine antibodies and PE-conjugated streptavidin. The fluorescence signal was detected using the Bio-Plex 200 system (Bio-Rad Inc., USA), and the cytokine concentration was estimated according to the standard curve prepared using the kit.

**Table 1:**
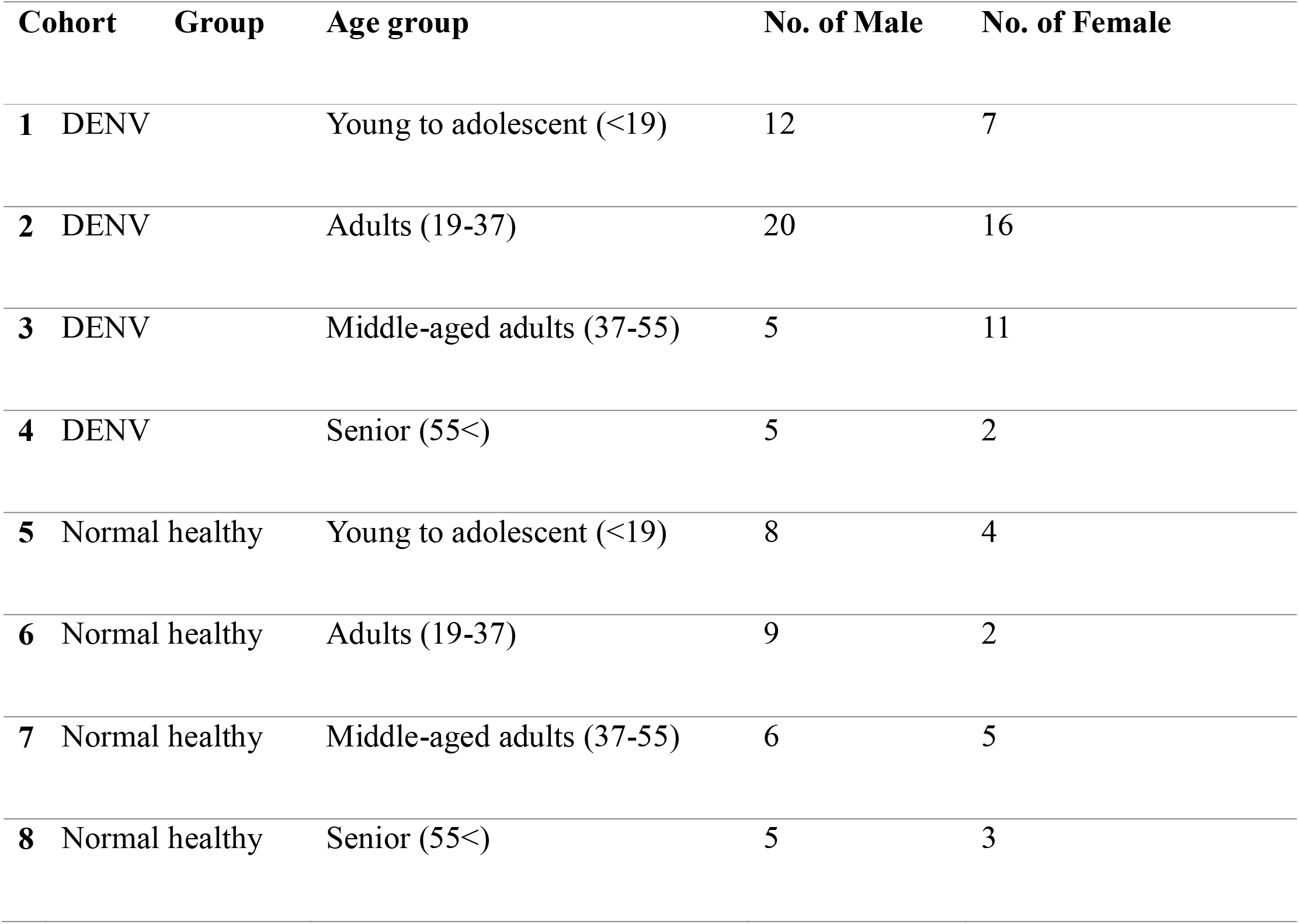
Different DENV and normal healthy serum samples analyzed based on the age and sex.

| **Cohort** | **Group** | **Age group** | **No. of Male** | **No. of Female** |
| --- | --- | --- | --- | --- |
| **1** | DENV | Young to adolescent (<19) | 12 | 7 |
| **2** | DENV | Adults (19-37) | 20 | 16 |
| **3** | DENV | Middle-aged adults (37-55) | 5 | 11 |
| **4** | DENV | Senior (55<) | 5 | 2 |
| **5** | Normal healthy | Young to adolescent (<19) | 8 | 4 |
| **6** | Normal healthy | Adults (19-37) | 9 | 2 |
| **7** | Normal healthy | Middle-aged adults (37-55) | 6 | 5 |
| **8** | Normal healthy | Senior (55<) | 5 | 3 |

## 3. Statistical analysis cum results

This study investigated seventy-eight patients with laboratory-confirmed dengue virus infection with the control groups for their cytokine profiles. We studied the levels of nine different cytokines: GM-CSF, IFN-γ, IL-2, IL-4, IL-5, IL-10, 1L-12 (P70), IL-13, and TNF-α in both the control and infected groups. The dataset is checked for normality distribution via different tests like the Anderson-Darling test, the D’Agostino & Pearson test, and the Shapiro-Wilk test, which showed the data deviates significantly from normality (Figure 1 (a)). However, few cytokines followed the normal distribution in one test but not in another. So, to conclude, we assumed none of the cytokines follow a normal distribution.

**Figure 1:**
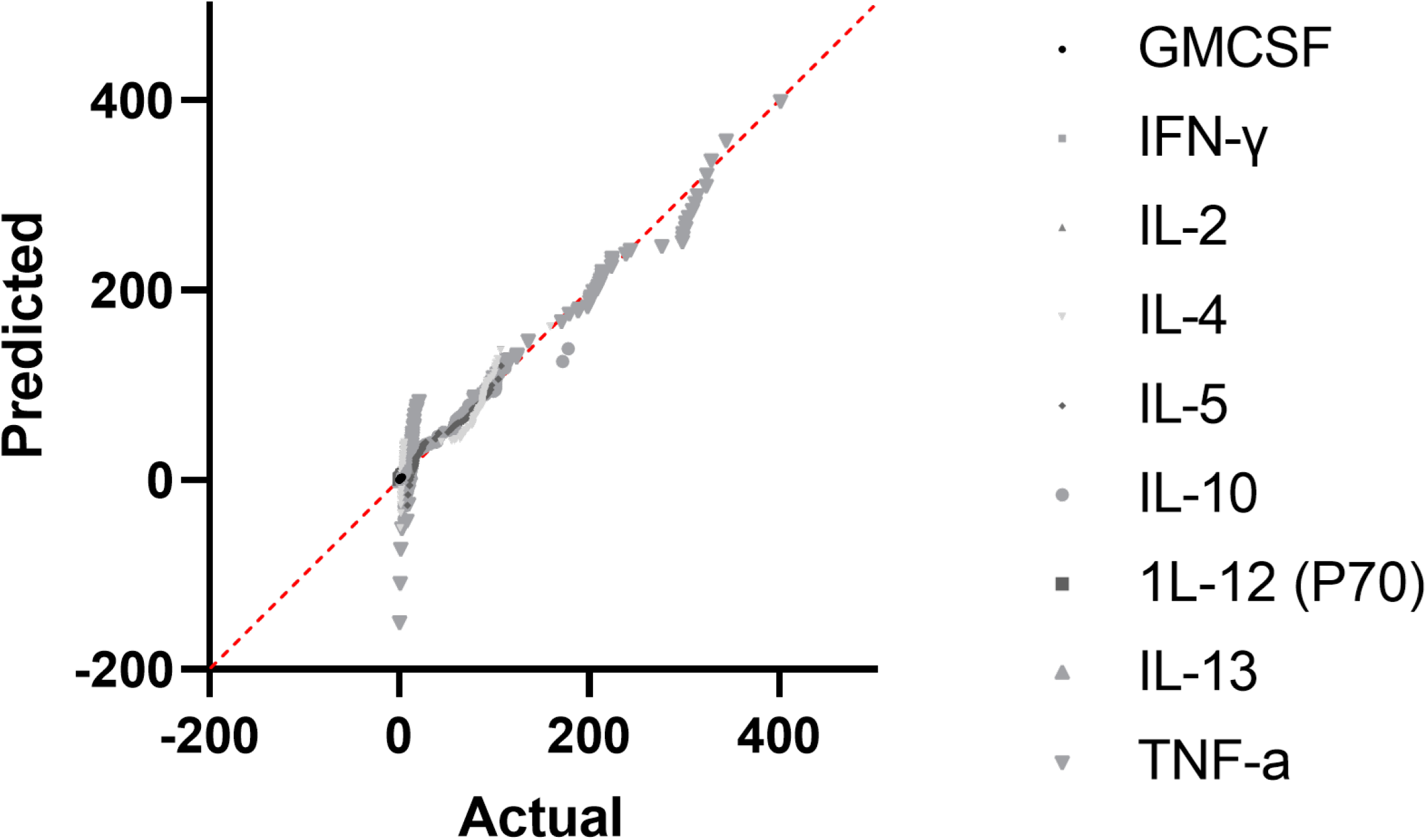
Expression of cytokines in normal healthy population.

### A. Relative levels of cytokines

On average, in the dataset, among all the cytokines, IFN-γ, IL-2, IL-4, IL-5, IL-10, IL-13, and TNF-α levels are consistently higher for patients than control across all age groups except GM-CSF, and 1L-12 (P70) (Figure 2 (a, g)). These two cytokines, GM-CSF and 1L-12 (P70) were found to be downregulated in the infected patients.

**Figure 2:**
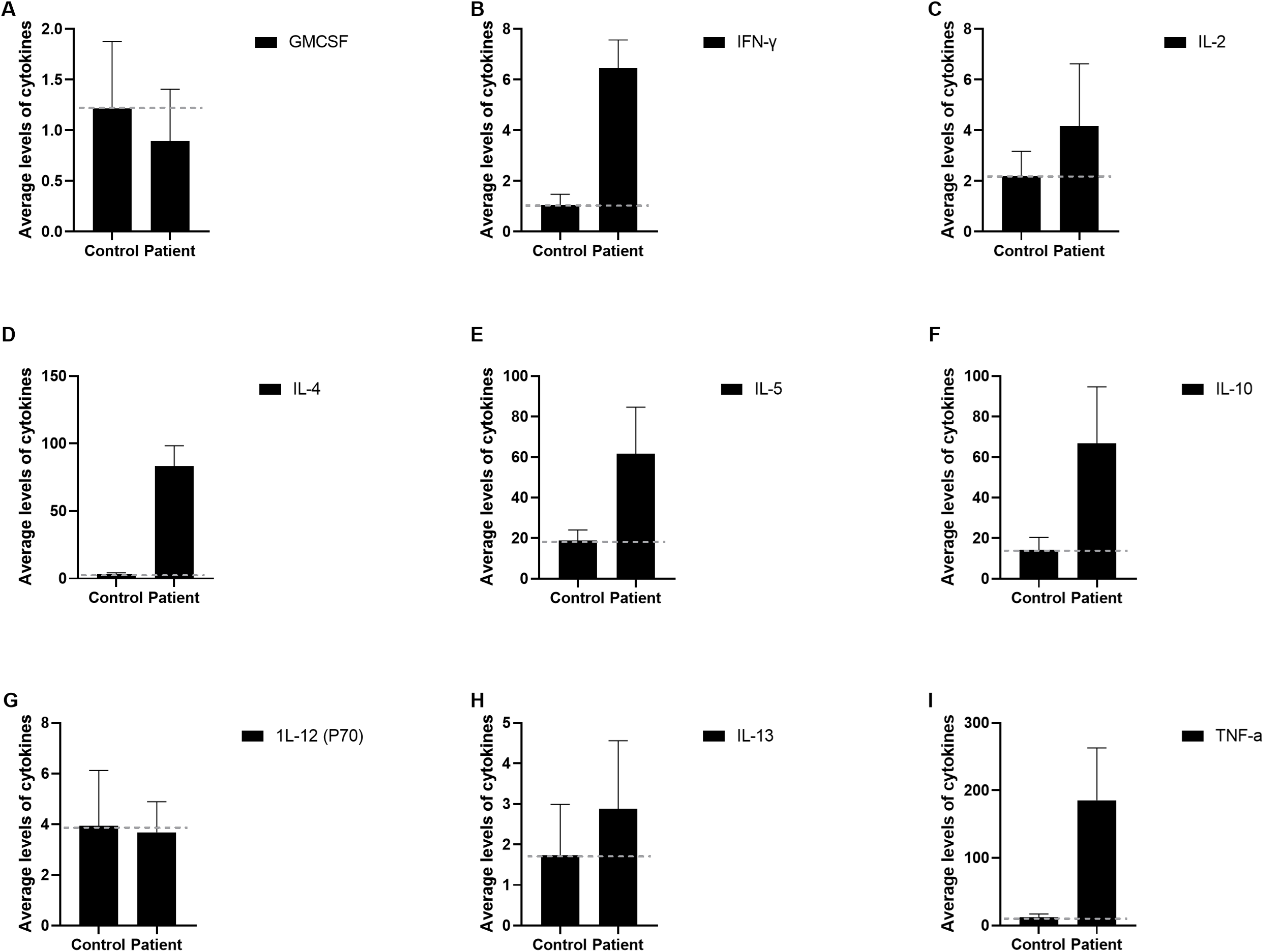
Levels of cytokines in the cohort of seventy-eight DENV-infected patients as compared to the control group. The grey line marks the threshold calculated based on the control group values.

Further analysis was carried out on the basis of age, sex, and difference in patient groups versus control groups (Figure 3). Females and males show varying correlations between cytokines (Figure 4). Notable differences were seen in the case of IL-4, IL-5, IL-10, and TNF-α. In females, IL-4, IL-5, and IL-10 levels were comparatively lower than in male patients, whereas TNF-α levels were higher for female patients. When age distribution is considered, IL-4, IL-5, IL-10, and TNF-α levels were found to differ like sex-based distribution (Figure 5). However, in the senior age groups, all the cytokines have a significantly higher level. Patients show an increasing trend of IL4 with age, peaking in the senior age (55<) group. However, when compared to the control group, values are relatively stable but slightly decline with age. IL5 peaks in the 55< age group, while the control group shows slight variations across groups. In the case of IL10 among infected individuals, it slightly decreases with age, but it remains consistently higher. TNFα shows a drastic difference between infected and control (Figure 2).

**Figure 3:**
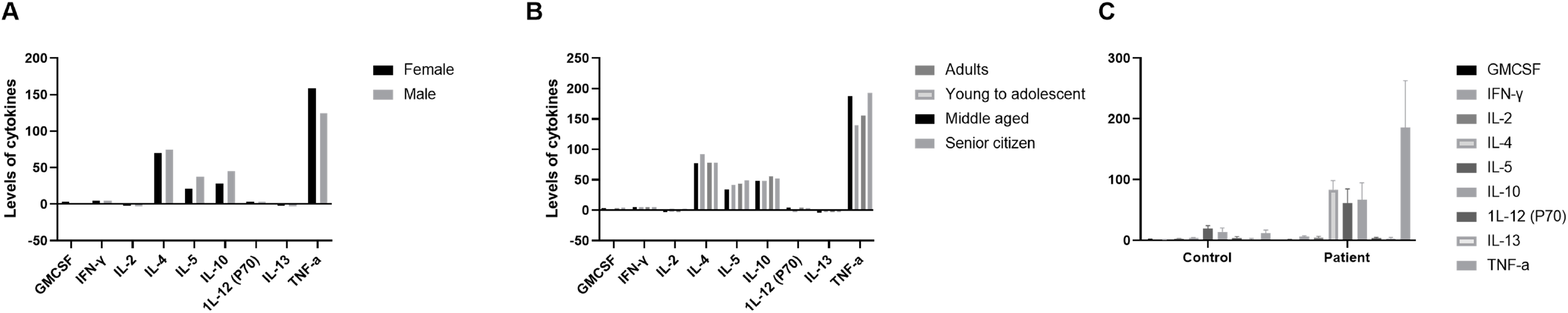
Difference in cytokines levels, keeping the control population as the threshold. (a) Comparative analysis based on the sex of the patient, (b) comparative analysis based on the age groups of the patient, and (c) Comparison between cytokine levels of control and patient.

**Figure 4:**
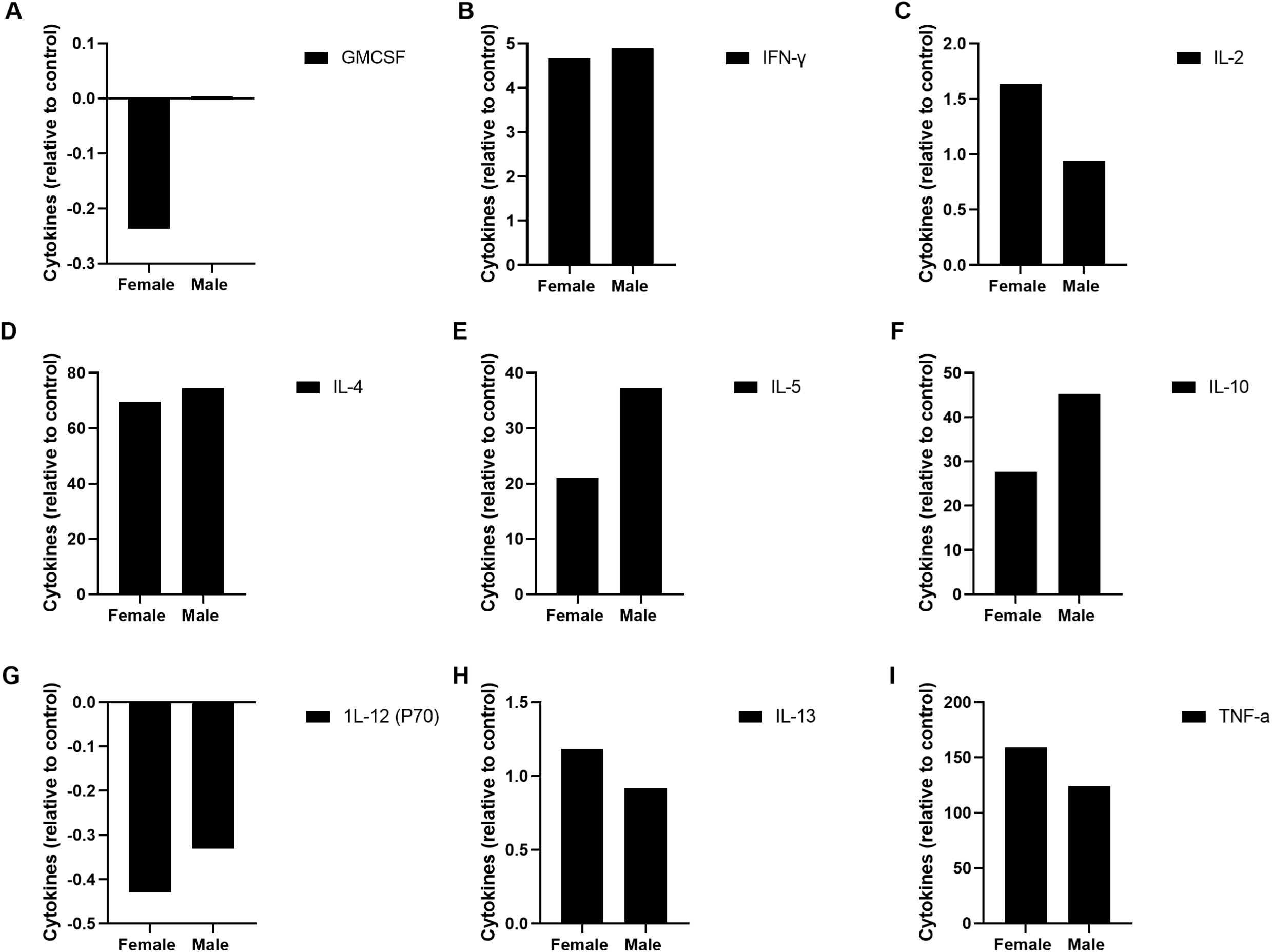
Difference of cytokine levels-based sex of the infected patients.

**Figure 5:**
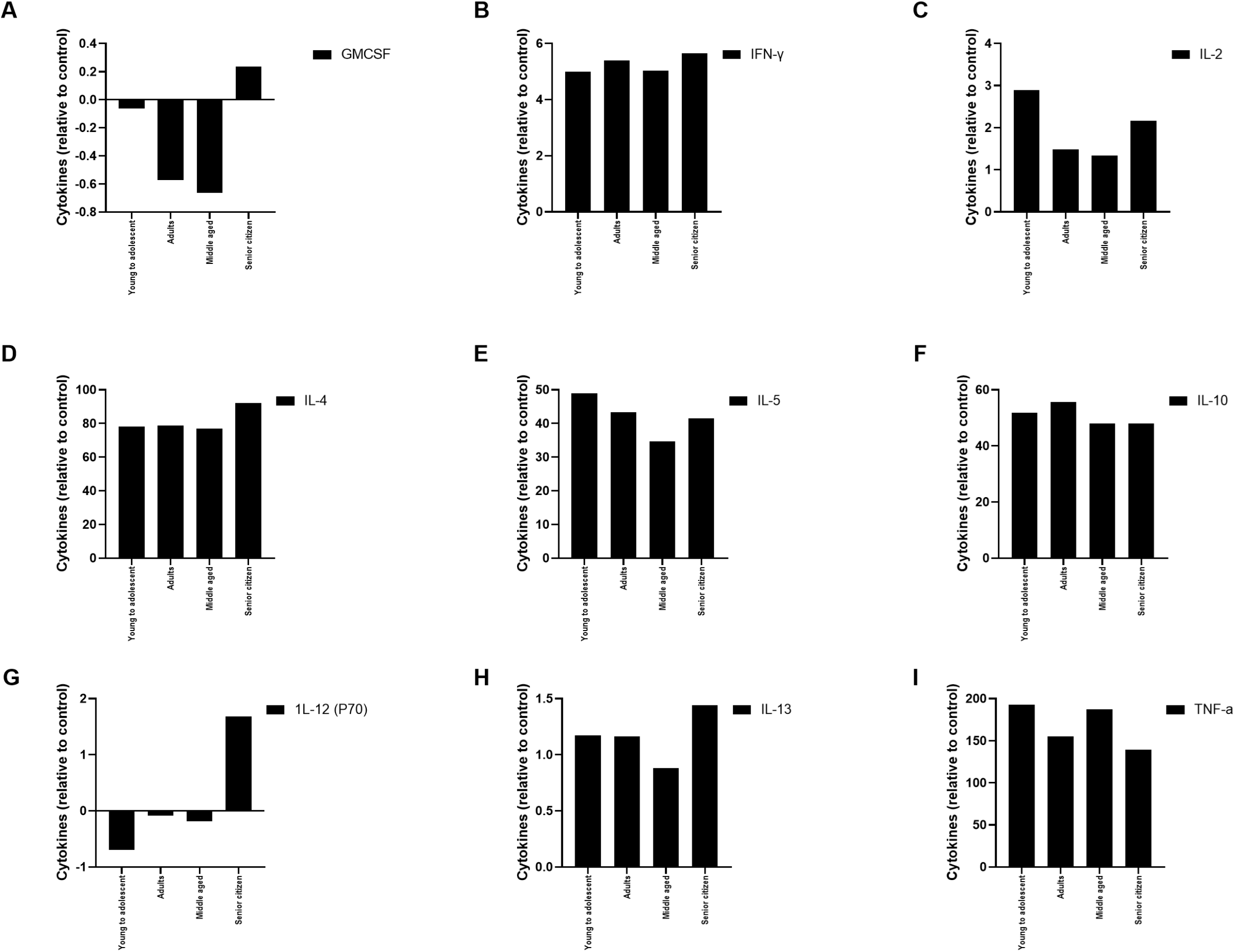
Comparison of cytokine levels across the age groups. The values are presented based on the control of the respective groups.

### B. Relationship between cytokine levels and demographics

Kruskal-Wallis one-way analysis of variance (ANOVA), followed by Dunn’s multiple comparison test, was used to evaluate differences between raw cytokine levels in the different groups of dengue patients as compared to control. However, the result failed to be analyzed based on sex and age, which could be due to many reasons; one could be less variability in the particular group. However, when the test was run between cytokines, significant differences were produced (Figure 6). The rank difference shows the size and direction of the difference; statistical significance indicates whether this difference is likely due to chance. For instance, “GM-CSF vs. IFN-γ” has a rank difference of -276.7. The negative sign means IFN-γ values are higher on average than GM-CSF, corresponding to their significant p-values.

**Figure 6:**
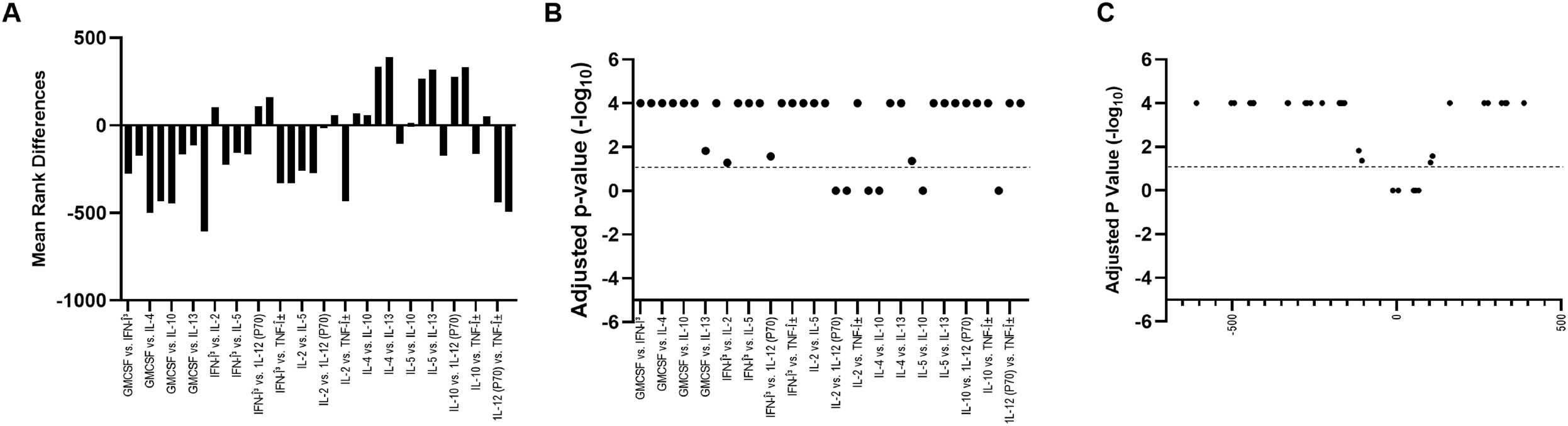
Kruskal-Wallis one-way analysis of variance results. (a) Mean rank difference between pairs of cytokines. (b) a p-value of the respective groups, and (c) a volcano plot of the adjusted p-values.

Also, the relationship between cytokine and grouping based on the sex and age of the patient and their interaction, a Multiple linear regression (MLR) test was carried out. MLR models the relationship between a dependent variable and two or more independent variables (predictors). Where significant obvious differences were noted for healthy versus infected individuals, the age group doesn’t produce any significant result, making it a weak predictor. Meanwhile, for the sex-based group, notable positive correlations were found for IL-5 and IL-10 (Figure 7). In the case of IL-5, the R-squared value comes to 0.586, indicating that 58.6% of the variance in IL-5 levels is explained by the model. Here, infected patients were strong predictors (obvious (p<0.001)), and sex was the moderate predictor for influencing the IL-5 levels (p=0.006p). Age and interaction group (sex and infected patient) showed no significant results. All predictors have VIF < 2, indicating no multicollinearity issues. The histogram suggests that residuals follow a normal distribution. No clear patterns, suggesting homoscedasticity (constant variance). Similar significant results were obtained for IL-10 (R-squared 0.58). The MLR model’s key findings indicate that both sex and infected patients significantly impact IL-5 and IL-10 levels, while age and the interaction term do not.

**Figure 7:**
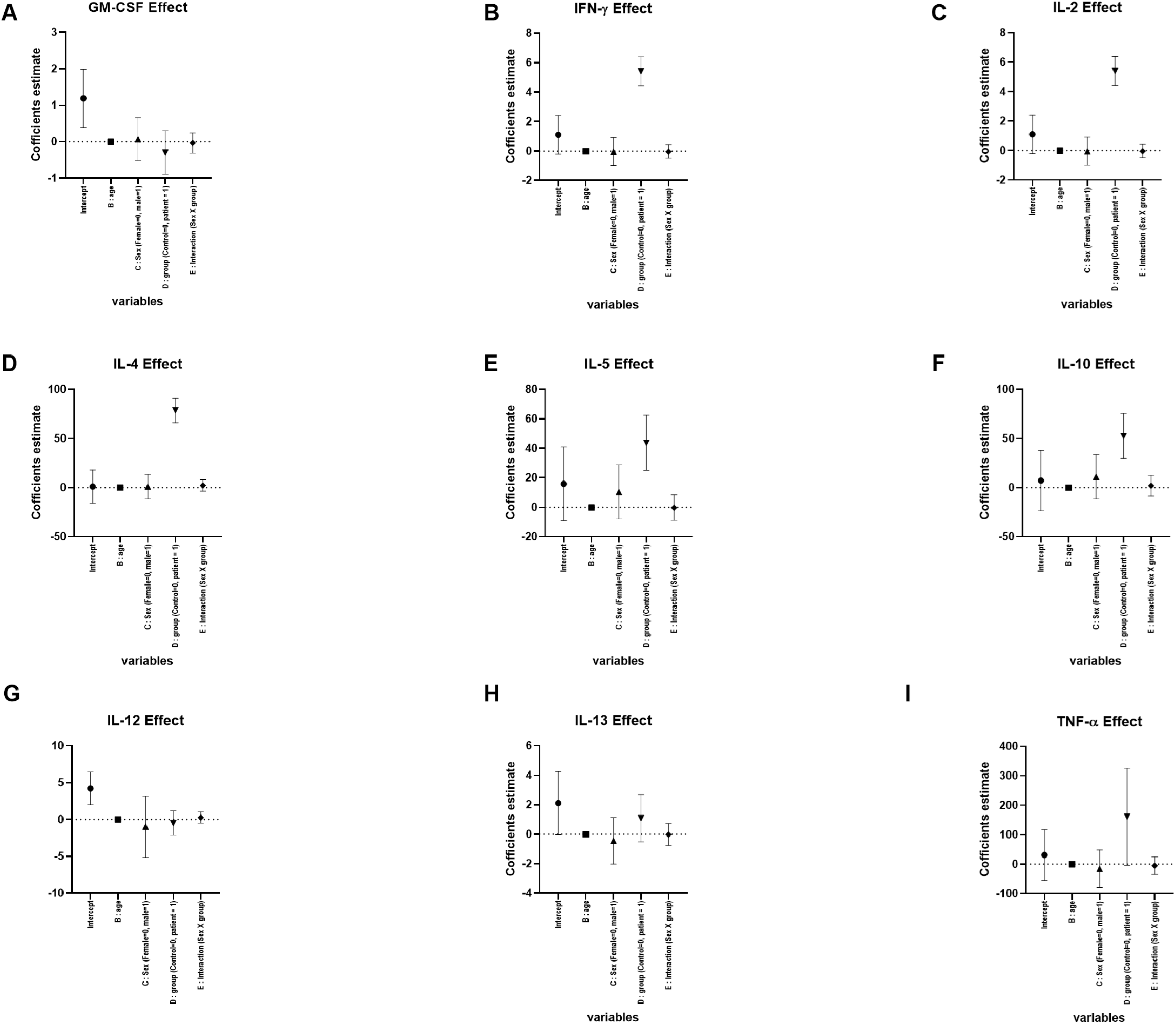
Multiple linear regression results (a-i). Here, the model has cytokine as the dependent variable and age, sex, group, and interaction (sex x group) as predictors.

## 4. Discussion

Analysis of cytokine levels from infected serum samples might help researchers and medical professionals understand the infection’s severity, the disease’s progression, and the complications that occur. Identifying cytokine patterns associated with different stages of dengue infection could lead to better diagnostic tools, risk stratification, and more effective treatment strategies. In dengue, the body’s immune response is complex, and an imbalance in cytokine levels can contribute to the severity of the disease. For instance, high levels of pro-inflammatory cytokines such as TNF-α, IL-6, and IL-10 have been associated with severe outcomes, while anti-inflammatory cytokines, like IL-4 and IL-10, may play a role in limiting the damage caused by the immune response. In the current study, IFN-γ, IL-2, IL-4, IL-5, IL-10, IL-13, and TNF-α cytokines are found to be higher in dengue-infected patients. Among these anti-inflammatory cytokines, IL-5 and IL-10 are shown to be regulated differently in male-female populations. However, it needs to be further investigated whether the predictors have established biological mechanisms explaining their role in influencing IL-5 and IL-10 levels. Eventually, cytokine profiling can improve our understanding of dengue virus pathogenesis and pinpoint specific immune regulators to manage the course of therapeutics in the area.

## 6. Acknowledgment

The virus research in the laboratory is supported by the Indian Council of Medical Research (IIRP-2023-2444).

